# Autonomous adaptive optimization of NMR experimental conditions for precise inference of minor conformational states of proteins based on chemical exchange saturation transfer

**DOI:** 10.1101/2024.10.07.616944

**Authors:** Takuma Kasai, Takanori Kigawa

## Abstract

In scientific experiments where measurement sensitivity is a major limiting factor, optimization of experimental conditions, such as parameters of measurement instruments, is essential to maximize the information obtained per unit time and the number of experiments performed. When optimization in advance is not possible because of limited prior knowledge of the system, autonomous, adaptive optimization must be implemented during the experiment. One approach to this involves sequential Bayesian optimal experimental design, which adopts mutual information as the utility function to be maximized. In this study, we applied this optimization method to the chemical exchange saturation transfer (CEST) experiment in nuclear magnetic resonance (NMR) spectroscopy, which is used to study minor but functionally important invisible states of certain molecules, such as proteins. Adaptive optimization was utilized because prior knowledge of minor states is limited. To this end, we developed an adaptive optimization system of ^15^N-CEST experimental conditions for proteins using Markov chain Monte Carlo (MCMC)to calculate the posterior distribution and utility function. To ensure the completion of MCMC computations within a reasonable period with sufficient precision, we developed a second-order approximation of the CEST forward model. Both simulations and actual measurements using the FF domain of the HYPA/FBP11 protein with the A39G mutation demonstrated that the adaptive method outperformed the conventional one in terms of estimation precision of minor-state parameters based on equal numbers of measurements. Because the algorithm used for the evaluation of the utility function is independent of the type of experiment, the proposed system can be applied to other instruments, as well as other NMR experiments, if the forward model or its approximation can be calculated sufficiently quickly.

**Author Summary:** In scientific experiments, experimental conditions are usually optimized to maximize the amount of information obtained about the target system. However, this optimization often involves measuring unknown system parameters. In nuclear magnetic resonance (NMR) spectroscopy, chemical exchange saturation transfer (CEST) is an effective method for observing minor states of certain molecules, such as proteins, indirectly via an easily observable major state. The optimal conditions for CEST depend on the resonant frequency of the minor state, which is not known before the experiment. Therefore, conventional CEST requires repeated measurements over a wide frequency range, most of which contain little information regarding minor populations. This problem may be resolved by adopting a step-by-step strategy that selects the most informative experimental condition for each subsequent step based on past data. In this study, we implemented this strategy for the CEST analysis of proteins. It can be also applied to other NMR experiments.

## 1. Introduction

One of the primary purposes of scientific experiments is to estimate parameters of mathematical models (henceforth referred to as model parameters) that represent or approximate systems of interest. Experimental design is optimized to satisfy specific statistical criteria, e.g., small variance in the estimated model parameters [1]. Bayesian design usually aims to maximize the amount of information gathered regarding the model and/or model parameters per unit cost. Thus, it adopts the expected gain in Shannon information in terms of the model parameters as the utility function to be optimized [2,3], which is equivalent to the expected Kullback-Leibler (KL) divergence as well as the mutual information [4–6]. Sequential or iterative design, which involves optimization based on recursive pairs of experimental and design phases, is expected to outperform pre-determined experimental designs in the case of nonlinear models as it leverages information obtained from previous observations [6–8]. On this basis, combined sequential and Bayesian design has also been suggested—however, this was difficult to be realized at the time of its proposal because of computational intractability [6]. Owing to methodological and hardware developments in recent years, the scope of this approach has expanded from simplified hypothetical problems to complicated real-world problems in various research fields [9–16].

To date, only a limited number of applications of sequential design have been reported in nuclear magnetic resonance (NMR) spectroscopy. High-Resolution Iterative Frequency Identification (HIFI) NMR was developed to estimate signal positions on original 3D spectra based on 2D projections via iterative selection of optimal projection angles [17]. Based on this method, an Assignment-directed Data collection Algorithm utilizing a Probabilistic Toolkit in NMR (ADAPT-NMR) was established to select informative angles for the stochastic assignment of protein NMR signals using a Bayesian network and belief propagation approach [18]. This approach calculated the utility function analytically to determine the optimal solution for networks containing loop structures [18]. However, it depended on ingenious model-specific treatments, thereby sacrificing generalizability. For simpler models with only two model parameters, Song et al. reported sequential optimization of experimental parameters for longitudinal or transverse relaxation measurements via exhaustive evaluation of the entire model parameter space [19]. Recently, an iterative selection method for evolution times and phases has been proposed in the context of non-uniform sampling for line-shape fittings. It utilizes linear approximation of a non-linear model [20]. Thus, the next step to expand the scope of NMR applications requires the development of sequential design methodologies capable of addressing relatively complicated nonlinear models in their intact forms, even at the cost of slightly higher computation loads.

Chemical shift exchange saturation transfer (CEST) is an NMR experiment that reveals minor and, therefore, directly invisible conformations of molecules, including proteins, slowly exchanging with a major visible state (Vallurupalli et al. [21] and references therein). Conventional CEST involving numerous sampling points corresponding to evenly spaced frequency offsets of saturation pulses is significantly time consuming [22,23]. To mitigate this problem, optimization of the offset step [23], interpolation of the sampling points in the frequency domain using linear prediction in the time domain [24], and multifrequency or cosine-modulated saturation pulses [25–27] have been proposed. In general, ^15^N-CEST experiments exhibit high signal-to-noise ratios (SNRs), because the typical protein concentration in ^15^N-CEST experiments is approximately 1 mM [21], with a few exceptions in the case of small peptides or relatively large (∼10 %) minor populations [28,29]. Therefore existing design approaches in the case of CEST assume sufficient sensitivity, thereby excluding repetition of the same experimental condition, analogous to the so-called “sampling-limited regime” in the context of non-uniform sampling or reduced dimensionality [30]. However, in the “sensitivity-limited regime,” repetitive sampling of important experimental conditions is essential to improve sensitivity via accumulation, leading to more precise model-parameter estimation. In the context of the preceding discussion, sequential Bayesian optimal design is a promising method because it samples informative experimental conditions preferentially.

In this study, a sequential Bayesian design method applicable to ^15^N-CEST experiments on proteins is proposed, assuming low SNR. To achieve this, we employed Markov chain Monte Carlo (MCMC) and Riemann sum approximations to calculate the utility function. As MCMC-based Bayesian analysis involves significantly more forward model evaluation times than point estimation analysis using a gradient-descent algorithm, we introduced a second-order approximation of the *R*_1ρ_ relaxation coefficient to accelerate the evaluation of the forward model. We compared the precision and accuracy of the proposed and conventional methods based on model parameters estimated via simulation as well as real observation of the FF domain of the HYPA/FBP11 protein with the A39G mutation.

## 2. Results and Discussion

### 2.1 Theoretical background of the proposed method

A detailed theoretical background of adaptive CEST is provided in Supporting Information. In this section, we present a brief summary. The pseudocode for adaptive CEST is as follows:

set the next experimental condition ^(1)^𝒙 to the reference (*ω*_RF_ = 0 Hz, *ω*_1_ = 0 Hz, 𝑇_EX_ = 0 s)

for iteration 𝑛,

perform CEST measurements with the condition ^(𝑛)^𝒙

process NMR data to obtain intensities ^(𝑛)^𝐘 = {^(𝑛)^𝑦 }^𝐾^_k=1_

sample from the posterior distribution *p*(𝚯|𝓓) via MCMC, where 𝓓 = {^(𝑛)^𝐘}^𝑁^_k=1_

calculate the utility function 𝑈(𝒙) using the MCMC samples

set the next experimental condition ^(𝑛+1)^𝒙, which maximizes 𝑈(𝒙)

repeat iteration 𝑁 times

if necessary, resample from the posterior distribution *p*(𝚯|𝓓) for detailed analysis The experimental condition 𝒙 comprised the offset, *ω*_RF_, the strength, *ω*_1_, and the duration, 𝑇_EX_. of the irradiation pulse. In general, a constant 𝑇_EX_ is adopted in conventional CEST experiments [22]. However, based on the similarity between CEST and *R*_1𝜌_ experiments [31], the adaptive adjustment of 𝑇_EX_ were considered in this study to improve the performance.

The model parameter, 𝚯, is a set of parameters corresponding to different residues, θ_𝑘_ = {*p_B_*^(*k*)^,*k_ex_*^(*k*)^,*ω_B_*^(*k*)^,*R_1_*^(*k*)^,*R_2A_*^(*k*)^,*R_2B_*^(*k*)^,𝐼*_0_*^(*k*)^}, where *p_B_* denotes the population ratio of the invisible (B) state, *k*_ex_ denotes the exchange rate, *ω*_B_ denotes the chemical shift of the B state, *R*_1_ denotes the longitudinal relaxation rate constant, *R*_2A_ denotes the transverse relaxation rate constant for the observable major (A) state, *R*_2B_ denotes that of the B state, and 𝐼_0_ denotes the basal intensity at 𝑇_EX_ = 0. The chemical shift of the A state, *ω*_A_, is known from the spectrum. *p*_B_ and *k*_ex_ are likely to be constant or correlated among residues in single-domain globular proteins. However, this is usually confirmed by performing local (individual) fitting before global fitting, which fixes *p*_B_ and *k*_ex_ for all residues. In adaptive CEST, *p*_B_ and *k*_ex_ are also conservatively assumed to be independent of residues at the experimental design stage, whereas, in the detailed analysis after the experiment, the global model may be used.

As the utility function, we selected mutual information, defined by 𝑈(𝒙) = 𝐼(*p*(**Ŷ**);*p*(𝚯)), which is commonly used in Bayesian design [5,6]. Here, **Ŷ** denotes the stochastic variable of the future observation, ^(𝑛+1)^**Y**. Mutual information is the expected value of the Kullback-Leibler (KL) divergence, *D*_KL_(*p*(𝚯|**Ŷ**)||*p*(𝚯)), over *p*(**Ŷ**) [4–6]. As we assumed independence of *p*(θ_*k*_) among residues, mutual information, 𝐼(*p*(**Ŷ**);*p*(𝚯)), was calculated as the sum of residues, 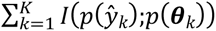, greatly reducing the parameter-space dimension from 7𝐾 to 7, thereby improving the efficiency of the MCMC. Note that the same 𝒙 can be repetitively selected corresponding to different iterations, which may be effective in low-SNR cases, analogous to the increase in the number of scans in conventional NMR.

### 2.2 Approximation of the CEST forward model with low computational cost

While performing autonomous measurements with a sequential Bayesian design, the posterior distribution and the utility function between the iterative measurements need to be evaluated within a short period to maximize the effectiveness of the NMR machine per unit time. Owing to the intrinsic similarities between CEST and *R*_1𝜌_ experiments, the CEST forward function has been approximated using *R*_1𝜌_ values in multiple papers, with reduced computational cost compared to that incurred in calculating the Bloch-McConnell equation completely [31,32]. While *R*_1𝜌_ can be calculated as one of the eigenvalues of the propagation matrix of the Bloch-McConnell equation, first-order approximations with some perturbations have been proposed to further reduce computational time [33,34]. However, as these approximations were not sufficiently accurate for our purpose, we approximated *R*_1𝜌_ to the second order (Supplementary Note S2, Supplementary Fig S1). When combined with the Palmer’s CEST approximation [31], the proposed *R*_1𝜌_ approximation method exhibited a typical computation time that exceeded those of first-order methods by only 25 %, and was lower than that required for numerical eigenvalue calculation by an approximate factor of 30 (Supplementary Table S1).

### 2.3 A simple 1-residue simulation

As a proof-of-concept for adaptive CEST, we first simulated a 200-iteration experiment, henceforth referred to as simulation A1, using a virtual 1-residue protein (Fig 1a). The experimental configurations and true model parameters are listed in Supporting Information (Supplementary Tables S2 and S4). The SNR, defined to be 𝐼_0_/𝜎, was taken to be 20, where 𝜎 denotes the standard deviation of the spectral background noise. The apparent SNR of the CEST curve was reduced to 12.2 when 𝑇_EX_ = 0.5 s because the off-resonance signal intensity was 𝐼_0_ exp( ― *R*_1_𝑇_EX_). There were 803 experimental condition candidates (𝒳), including the reference (𝑇_EX_ = 0 s) for the simulation A1. The offset *ω*_RF_spanned from -1000 to 1000 Hz, and the strength *ω*_1_ was either 10 or 50 Hz.

**Fig 1.**
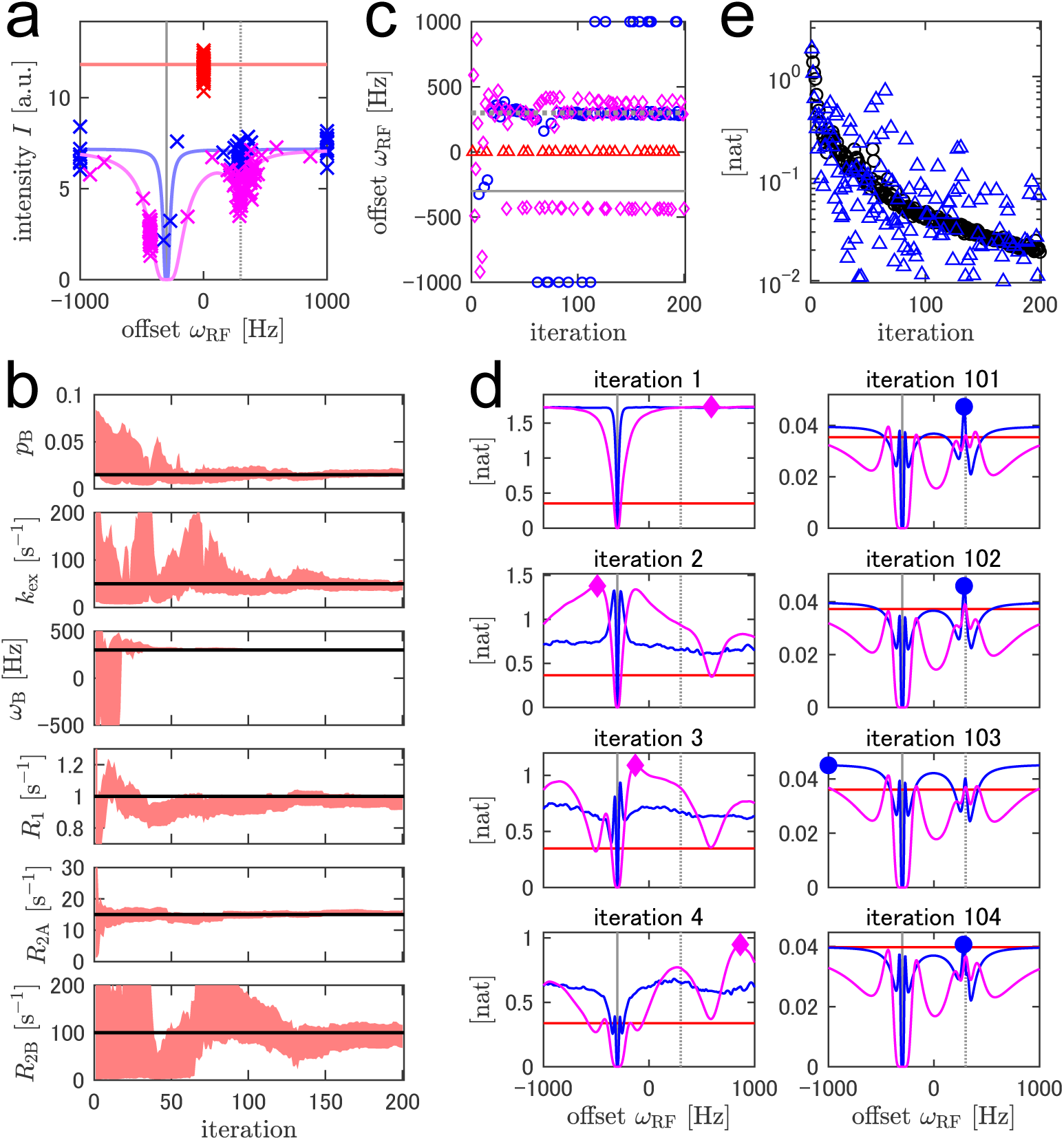
Simulated adaptive ^15^N-CEST with a single-residue virtual protein. (a) Observed signal intensities in all 200 iterations are indicated by crosses. The theoretical noiseless responses are indicated using lines. The colors represent different values of the irradiation strength, *ω*_1_: 0 Hz = red, 10 Hz = blue, and 50 Hz = magenta. Vertical solid and dotted gray lines represent the chemical shifts of A (*ω*_𝐴_) and B (*ω*_𝐵_) states, respectively. (b) Red areas represent 68.3 % credible intervals (CIs) for the estimated model parameters. Horizontal black lines represent the actual parameter values. (c) The selected irradiation offsets, *ω*_*R*𝐹_, are plotted indicating different values of *ω*_1_ using different markers: 0 Hz = red triangle, 10 Hz = blue circle, and 50 Hz = magenta diamond. (d) The mutual information evaluated following observations recorded at the designated representative iterations. The experimental condition with the highest mutual information is indicated—it is selected for the next iteration. The same markers as those used in (c) were used. *ω*_𝐴_ and *ω*_𝐵_ are indicated by vertical lines, as in (a). (e) The mutual information (black circle) and KL divergence calculated based on the realized observation (blue triangle).

Fig 1b illustrates the 68 % CIs of the model parameters plotted with respect to varying numbers of iterations. Unsurprisingly, the estimation of model parameters became more precise as the number of iterations was increased. During the first 20 iterations, there was almost no information regarding the chemical shift of the B state. After that, it was confirmed that *ω*_B_ was ∼300 Hz, and prompting us to gather information about other exchange parameters (*p*_B_, *k*_ex_, *R*_1_, *R*_2A_, and *R*_2B_). This change also appeared during the selection of the experimental conditions. In the former iterations, the selected offset values were varied to search for a B-state dip, whereas in the later iterations, the offset values were more stable (Fig 1c). The selection of the experimental conditions during the later iterations was qualitatively explained using the simulated CEST curves. Slight changes in *p*_B_ and *k*_ex_ values affected the dip size of the B state (Supplementary Fig S3). This indicated that the B-state on-resonance experiments (*ω*_RF_≅*ω*_B_ = 300 Hz) were selected because they were informative for *p*_B_ and *k*_ex_. Similarly, the slight off-resonance corresponding to the B-state (*ω*_RF_≅400 Hz) experiments was for *R*_2B_; the slight off-resonance corresponding to the A-state (*ω*_RF_≅ ― 450 Hz) experiments was for *R*_2A_; the off-resonances corresponding to *ω*_RF_ = ―1000 or 1000 Hz) was for *R*_1_, and the reference experiments were for 𝐼_0_. However, because some of the model parameters were correlated, information on one parameter aided the estimation of others (Fig S4).

Fig 1d depicts the mutual information as the utility function for representative iterations. For the first four iterations, the shape of the function varied drastically, suggesting that the posterior distributions, especially of *ω*_B_, were considerably updated by informative observations (Fig 1d: left panel). Consequently, various offset values were selected to search for the B state dip during these iterations (Fig 1c). In contrast, after reaching confidence for *ω*_B_, the shape of the function was stable with *ω*_RF_ being restricted to the aforementioned typical conditions (∼300 Hz, ∼400 Hz, ∼−450 Hz, off resonances, or the reference) at these iterations (Fig 1d right). As discussed in the Theory section, mutual information is expected to exhibit KL divergence over the observation distribution, *p*(**Ŷ**). Fig 1d depicts the mutual information evaluated prior to observation and the KL divergence evaluated using the realized **Ŷ** after observation. The mutual information decreased as the number of iterations increased (Fig 1e). This was attributed to a general lack of new information obtained from a single future observation compared to the knowledge pieced together based on numerous past observations as the number of iterations increased. In the first few iterations, the probability density of *p*(**Ŷ**) was widespread, indicating the difficulty of prediction (Fig 2a-c). In such cases, the system confirmed unpredictable observations, as explained in the literature, where the observations were considered to be virtually free from noise [35]. However, this explanation no longer held true after the prediction of uncertainty was reduced to the noise level. In this case, the smaller KL divergence compared to that of mutual information reduced the difference between the posterior distributions before and after the observation. For example, during Iteration 32, 68% CI of the intensity of the subsequent observation was predicted to be 2.06–3.37 (Fig 2d). Because the observation at the Iteration 33fell within the predicted range (2.73), the KL divergence was as small as 0.0167 nat, which was lower than the mutual information of 0.134 nat (Fig 2d). Such well-predicted measurements increased confidence in current knowledge. In contrast, KL divergences larger than the mutual information indicated an outlying observation derived from significant noise and/or inaccurate prediction. The former case was instantiated in Iteration 83, which featured large KL divergence owing to accidental large noise, even though the prediction of the observation was close to the true distribution (Fig 2f). The latter case was instantiated in Iteration 35. The 68 % CI of the subsequent observation was 4.95–6.32, whereas the true value was 4.46. The disparity between the predicted range and the observation led to a larger KL divergence of 0.293 nat compared to the mutual information of 0.101 nat, as the observed divergence was significantly smaller than the predicted divergence (Fig 2e). Such informative observations increase the agreement between the knowledge gathered and the true values of the model parameters. As mutual information is defined as the expected value of KL divergence, 𝐼(*p*(**Ŷ**);*p*(𝚯)) = 𝔼_**Ŷ**_[*D*_KL_(*p*(𝚯|**Ŷ**)||*p*(𝚯))] = ∫ *p*(**Ŷ**)*D*_KL_(*p*(𝚯|**Ŷ**)||*p*(𝚯))𝑑**Ŷ**, the possibility of large knowledge multiplied with its probability updates contributes the mutual information. In other words, the system autonomously designs experiments to decrease the risk of misevaluating the model parameters.

**Fig 2.**
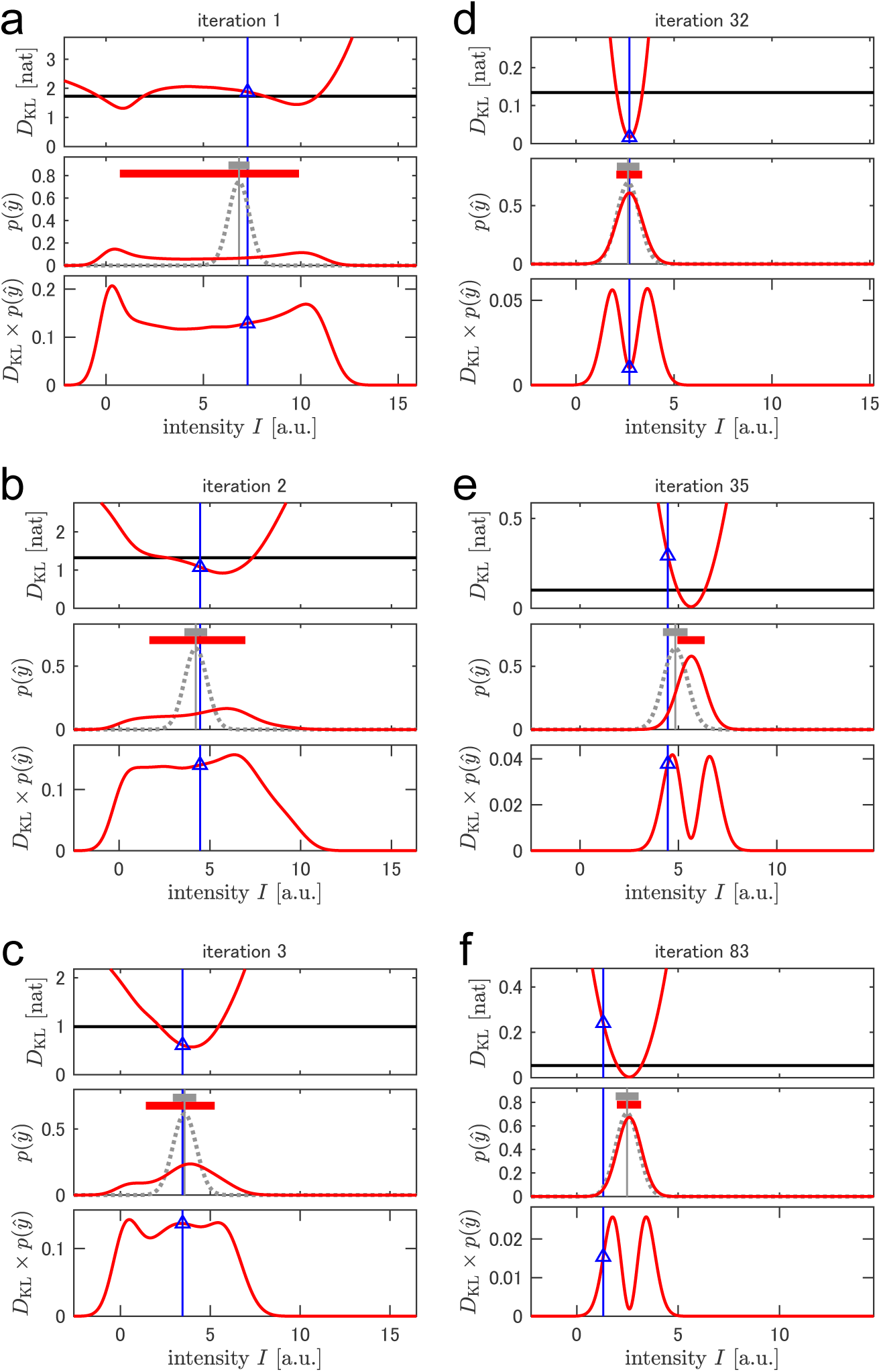
A breakdown of mutual information as the expected KL divergence and its difference with the realized KL divergence in single-signal simulation. In each panel, the mutual information at the designated iteration and the KL divergence calculated after the observation at the following iteration are depicted. The red line at the top represents the KL divergence, *D*_𝐾𝐿_(*p*(𝜃|*ŷ*)||*p*(𝜃)), plotted with respect to the future observation, *ŷ*. The red line in the middle indicates the estimated *p*(*ŷ*) calculated based on the current knowledge plotted against *ŷ*. The gray vertical line indicates the ideal noiseless response calculated using the true parameters. The gray dotted line represents the true distribution with a random observation noise. The red and gray horizontal thick lines represent the 68.3 % CI of the estimated and the true *p*(*ŷ*), respectively. The product of the above two functions of *ŷ*, *D*_𝐾𝐿_(*p*(𝜃|*ŷ*)||*p*(𝜃)) × *p*(*ŷ*), is represented by the red line at the bottom. As KL divergence estimates the degree of knowledge update after each subsequent observation, it tends to increase significantly when the subsequent observation is unexpected. Over the first iterations (i.e., at Iteration 1, 2, and 3), the knowledge about the model parameters is limited; thus, the estimated range of *p*(*ŷ*) is wide. In contrast, at later iterations (e.g., at Iteration 32, 35, and 83), the range of the estimated *p*(*ŷ*) reaches a width equal to the observation noise. As a result, the product, *D*_𝐾𝐿_(*p*(𝜃|*ŷ*)||*p*(𝜃)) × *p*(*ŷ*), exhibits trapezoidal or bimodal shapes in the former and latter cases, respectively. Mutual information, represented here by a black horizontal line, is the integration of the product, ∫ *D*_𝐾𝐿_(*p*(𝜃|*ŷ*)||*p*(𝜃))*p*(*ŷ*)𝑑*ŷ*. Blue vertical lines in all plots indicate the realized observation at each successive iteration. The blue triangles in the top and the bottom panels represent the corresponding realized *D*_𝐾𝐿_(*p*(𝜃|*ŷ*)||*p*(𝜃)) and *D*_𝐾𝐿_(*p*(𝜃|*ŷ*)||*p*(𝜃)) × *p*(*ŷ*), respectively.

### 2.4 Simulation with different irradiation durations

In conventional CEST, irradiation pulses are usually applied with various offsets, *ω*_RF_, a single or several strengths, *ω*_1_, and a fixed duration, 𝑇_EX_ [22]. This simple experimental configuration is beneficial to both experimental design as well as analysis of the results. However, we varied 𝑇_EX_ to improve parameter estimations, motivated by the close resemblance between CEST and *R*_1𝜌_ experiments, where 𝑇_EX_ is varied while investigating *R*_1𝜌_ via curve fitting [31]. Because the design and analysis steps of the adaptive experiment are free from human instruction, increasing the dimensionality of the design space from two to three is acceptable.

To exemplify incorporating 𝑇_EX_ as an experimental parameter, we tested a single-residue simulation named A2 with the same configuration as the simulation A1, except for fixed *ω*_1_ and variable values of 𝑇_EX_ over 0.50, 0.75, and 1.00 s (Supplementary Table S2). The model-parameter estimation was less precise than that for A1 (Supplementary Fig S5), possibly due to the lack of variance of *ω*_1_ [21,24]. For precise estimation of relaxation constants such as *R*_1𝜌_, 𝑇_EX_ = ^1^/*R*_1𝜌_ was observed to be the most informative sampling point when 𝐼_0_ was known. The selected value of 𝑇_EX_ was close to the inverse of the estimated 1/*R*_1𝜌_, with some exceptions possibly derived from the uncertainty of *R*_1𝜌_ and 𝐼_0_ estimations (Fig 3a). When the *R*_1𝜌_ estimate was large, smaller values of 𝑇_EX_ increased reliability. However, the approximation of the CEST forward model used in this study assumed sufficiently long 𝑇_EX_ to eliminate transverse magnetization in the tilted reference frame [31]. This was also true in case of the Bloch-McConnell equation because the complete measurement of 𝐵_1_ inhomogeneity is difficult [36]. For these reasons, we set the lower limit of 𝑇_EX_ as 0.5 s to ensure the accuracy of forward calculation. The curves of the utility function corresponding to different 𝑇_EX_ were similar even in the first iterations (Fig 3b), in contrast to those corresponding to different *ω*_1_ in the simulation A1 (Fig 1e). Therefore, we decided to use only 𝑇_EX_ = 0.5 or 1.0 s as candidates for the following simulations and measurements, conserving computational resources for the more important *ω*_1_ variation.

**Fig 3.**
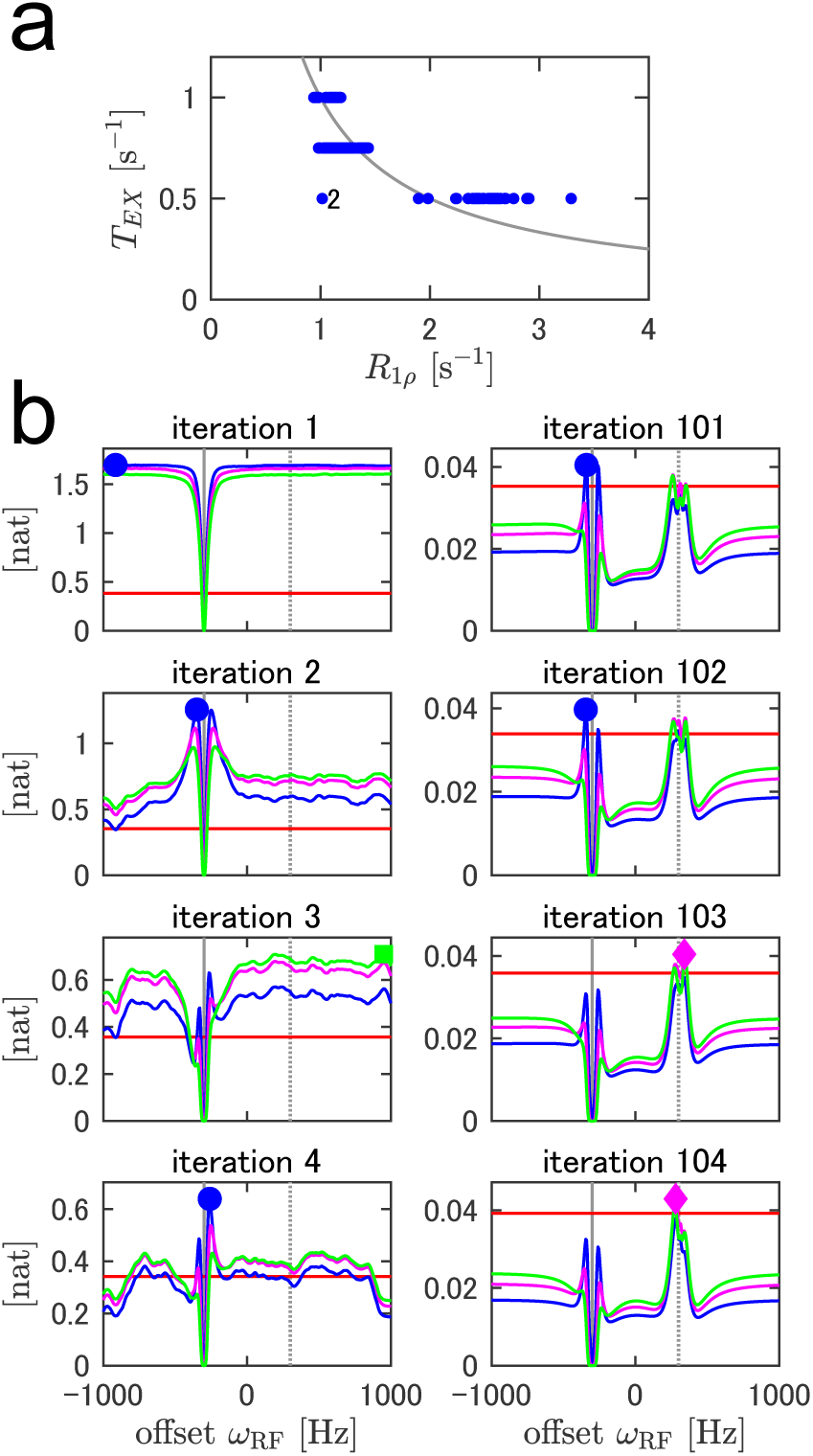
Adaptive CEST simulation of the virtual single-residue protein with variable irradiation durations, 𝑇_𝐸𝑋_. (a) Selected 𝑇_𝐸𝑋_ values with respect to maximum-a-posteriori *R*_1𝜌_. The gray line represents 𝑇_𝐸𝑋_ = 1*R*_1𝜌_. An outlier is observed at iteration 2, indicated by the letter “2”. (b) Mutual information at several representative iterations. The red, blue, magenta, and green lines correspond to 𝑇_𝐸𝑋_ = 0 𝑠 (reference), 0.50 𝑠, 0.75 𝑠, and 1.00 𝑠, respectively. The different markers represent selected experimental conditions with the highest mutual information.

### 2.5 Simulation involving a virtual 70-signal protein

In practical ^15^N-CEST experiments on proteins, we observed multiple signals corresponding to all residues, except prolines and an N-terminal residue. We defined the utility function to be the mutual information in the 7𝐾-dimensional model parameter space, i.e., the sum of the mutual information of individual 𝐾 signals (see the Theory section for details). In this situation, because the same experimental conditions were applied to signals with potentially different model parameters at different iterations, the conditions may not have been optimal for each signal. To evaluate the performance of multiple signals, we simulated adaptive (A3 and A4) and conventional (C4) CEST using a virtual 70-signal protein (Supplementary Table S2). The number of 2D measurements was set to 192 for all simulations to ensure a fair comparison of their performances based on equal instrumental resources. For adaptive CEST, *ω*_1_ candidates were either 6.3, 13.0, 26.2, and 50 Hz (for A3) or 6.3, 13.0, and 26.2 Hz (for A4). For conventional CEST, *ω*_RF_ values comprised 63 evenly spaced points between -1000 and 1000 Hz (32.3-Hz step); *ω*_1_ values were the same as in A4; and 𝑇_EX_ was fixed to 0.5 s. Including the three references (*ω*_1_ = 0 Hz, 𝑇_EX_ = 0 s), 63 × 3 + 3 = 192 2D measurements were performed in aggregate. All simulations were repeated ten times with different random seeds. Both conventional and adaptive CEST data were subjected to Bayesian analysis to compare the uncertainties of model parameters. Both the precision and accuracy of model parameter estimation for adaptive CEST were better than or comparable to those for conventional CEST corresponding to most signals (Fig 4a). Notably, the lower bound of the CIs of *R*_2B_ was higher in adaptive CEST. i.e., more information was obtained about *R*_2B_.

**Fig 4.**
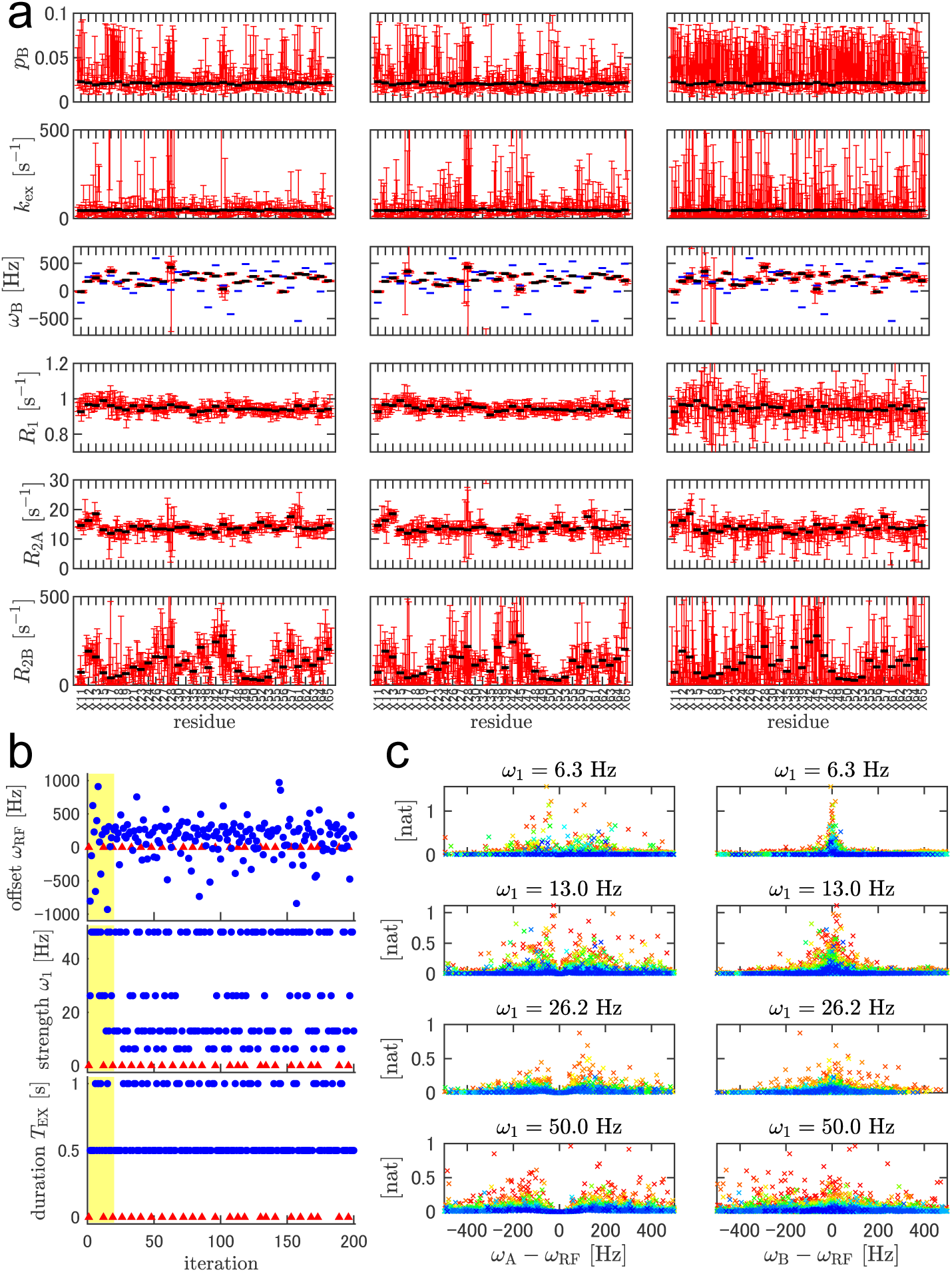
^15^N-CEST simulations with a virtual 70-signal protein. (a) Model parameter estimation after 192 iterations or 192 2D measurements. The left, middle, and right panels correspond to adaptive CEST with *ω*_1_ = 6.3, 13.0, 26.2, and 50.0 Hz; adaptive CEST with *ω*_1_ = 6.3, 13.0, and 26.2 Hz; and a conventional CEST with *ω*_1_ = 6.3, 13.0, and 26.2 Hz; respectively. The red lines represent the 68.3% CIs of the model parameter estimates. For each residue, 10 individual simulations with different random seeds are performed and plotted. The horizontal black lines represent the true values of the parameters. The horizontal blue lines in the *ω*_𝐵_ plots represent *ω*_𝐴_. Terminal residues and residues with |*ω*_𝐵_ ― *ω*_𝐴_| < 100 𝐻𝑧 are omitted. Plots with all residues are included in Supporting Information. (b) The selected experimental condition for adaptive CEST simulation with *ω*_1_ = 6.3, 13.0, 26.2, and 50.0 Hz. Blue circles and red triangles correspond to non-reference and reference experiments, respectively. The first 20 iterations are highlighted in yellow. (c) Mutual information of adaptive CEST simulation with *ω*_1_ = 6.3, 13.0, 26.2, and 50.0 Hz plotted against *ω*_𝐴_ ― *ω*_*R*𝐹_ and *ω*_𝐵_ ― *ω*_*R*𝐹_. Only Iterations 21–192 are illustrated, colored by a spectrum ranging from red (at Iteration 21) to blue (at Iteration 192).

Owing to variations in *ω*_A_, *ω*_B_, and the other model parameters among the signals, the optimum experimental condition depended on the signal. However, adaptive CEST optimized the experimental design notwithstanding the variety of optima. As in the case of the 1-signal simulation A1, the selected experimental condition for A3 over the first ∼20 iterations was varied to investigate B-state chemical shifts (Fig 4b). During this stage, strong irradiation (*ω*_1_ = 50 Hz) was preferred for effective exploration with a wide irradiation range. Over the rest of the iterations, both strong (*ω*_1_ = 50 Hz) and weak (*ω*_1_ = 6.3 or 13.0 Hz) strengths were selected. The mutual information of the individual residues plotted against *ω*_A_ ― *ω*_RF_ or *ω*_B_ ― *ω*_RF_ indicated that stronger irradiation facilitated the collection of information from a wide range of signals while weaker irradiation facilitated the collection of more detailed information corresponding to fewer signals (Fig 4c). Adaptive CEST was autonomously balanced using these two types of irradiation by following the definition of the utility function. It should be noted that the use of both strong and weak irradiation is important for adaptive CEST for this reason, together with the reported importance of exchange parameter estimation in conventional CEST [21,24]. In contrast to the remaining residues, A3 and A4 underperformed corresponding to residue X28 (Fig 4a), possibly owing to its outlying *ω*_B_ value. The proposed adaptive CEST autonomously prioritized estimation for the majority of signals, rather than for a single outlying signal based on the definition of the utility function. To focus on a specific signal, a different definition of the utility function should be used, e.g., residue-specifically weighted mutual information. Moreover, additional iterations with different utility functions would be useful for gathering more information about a specific residue after the experiment. Simulation A3, followed by an additional 48 iterations using the new utility function, i.e., the mutual information of residue X28, outperformed the simulation with the same number of iterations using the same utility function in terms of parameter estimation for residue X28 (Supplementary Fig S6).

### 2.6 Evaluation using Real Measurements

Finally, we performed a real CEST experiment with the 71-aa FF domain the HYPA/FBP11 protein with A39G mutation [22] (Fig 5a) using the same experimental configurations as those used in simulations A3 and C4. For adaptive CEST, MCMC and mutual information calculations were performed on a remote computer, which transmitted messages of selected experimental conditions to an NMR-control computer to run iterations (Fig 6, see experimental section for details). The measurement time for a single experimental condition, i.e., a single 2D spectrum, was 19 and 24 min for 𝑇_EX_ = 0.5 and 1.0 s, respectively. The typical computation times for MCMC and mutual information were 50–60 s and 18 s, respectively, using dual Intel Xeon E5-2690v4 CPUs (a total of 28 physical cores). In combination with the communication time, the typical turnaround time between NMR measurements was less than 90 s, which slightly increased the total occupation time of the machine.

**Fig 5.**
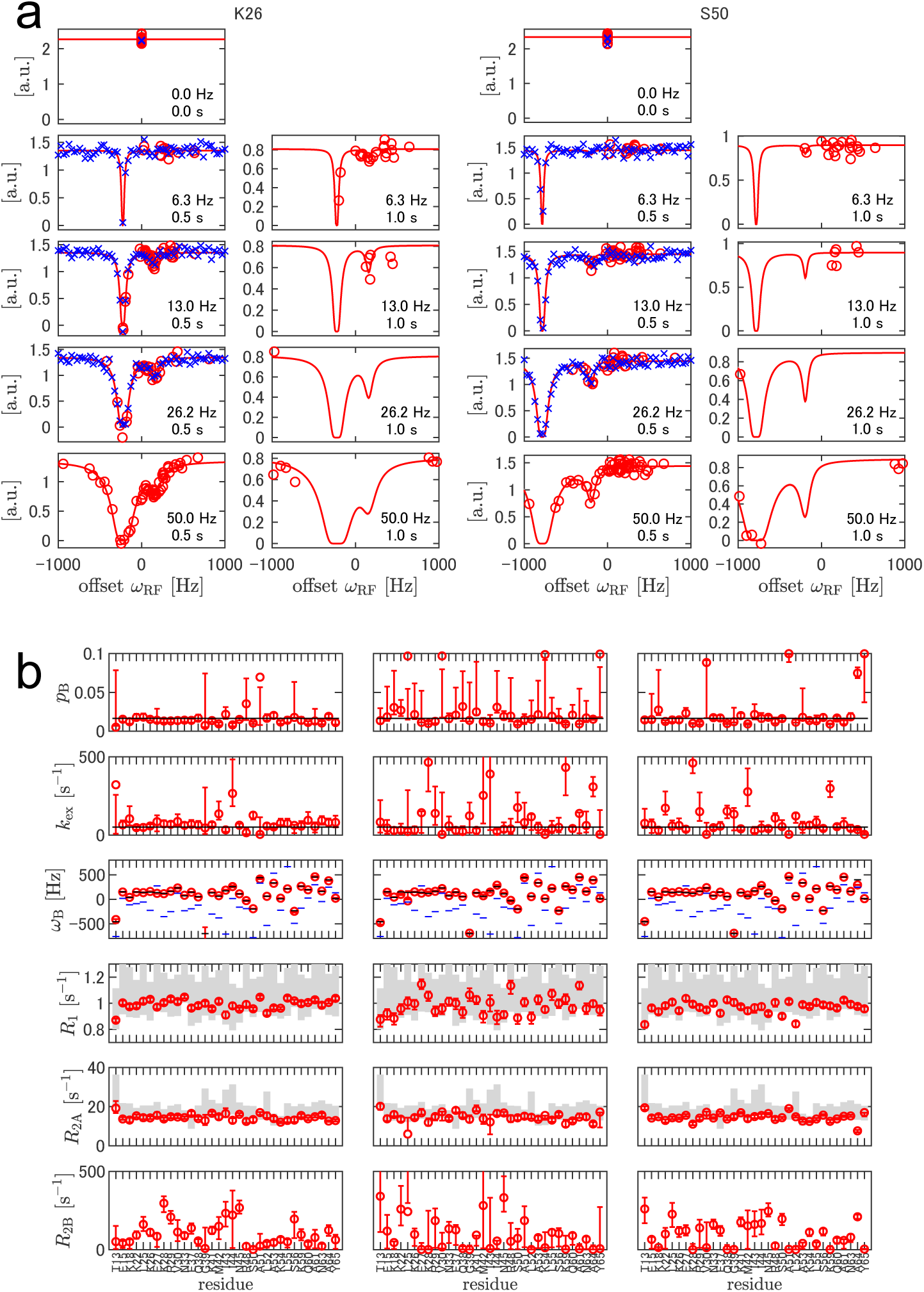
Adaptive ^15^N-CEST experiment with a 0.1 mM FF A39G protein, compared with conventional experiments. (a) Observed intensities of the two representative residues in the adaptive experiment (red circle) and the conventional experiment (blue cross). *ω*_1_ and 𝑇_𝐸𝑋_values are indicated in each plot. Red curves represent the responses calculated using the MAP estimator after 192 iterations of the adaptive experiment. (b) Estimated model parameters of adaptive CEST with 192 iterations (left), conventional CEST with 192 2D measurements (middle), and conventional CEST with 768 2D measurements (right). The red circles and error bars represent the MAP estimator and 68.3 % CIs, respectively. The black horizontal lines represent the reported parameters from conventional CEST with 2.0 mM protein [22]. The gray bars indicate 68.3 % CIs from the separately recorded ^15^N relaxation experiments with the same 0.1 mM sample.

**Fig 6.**
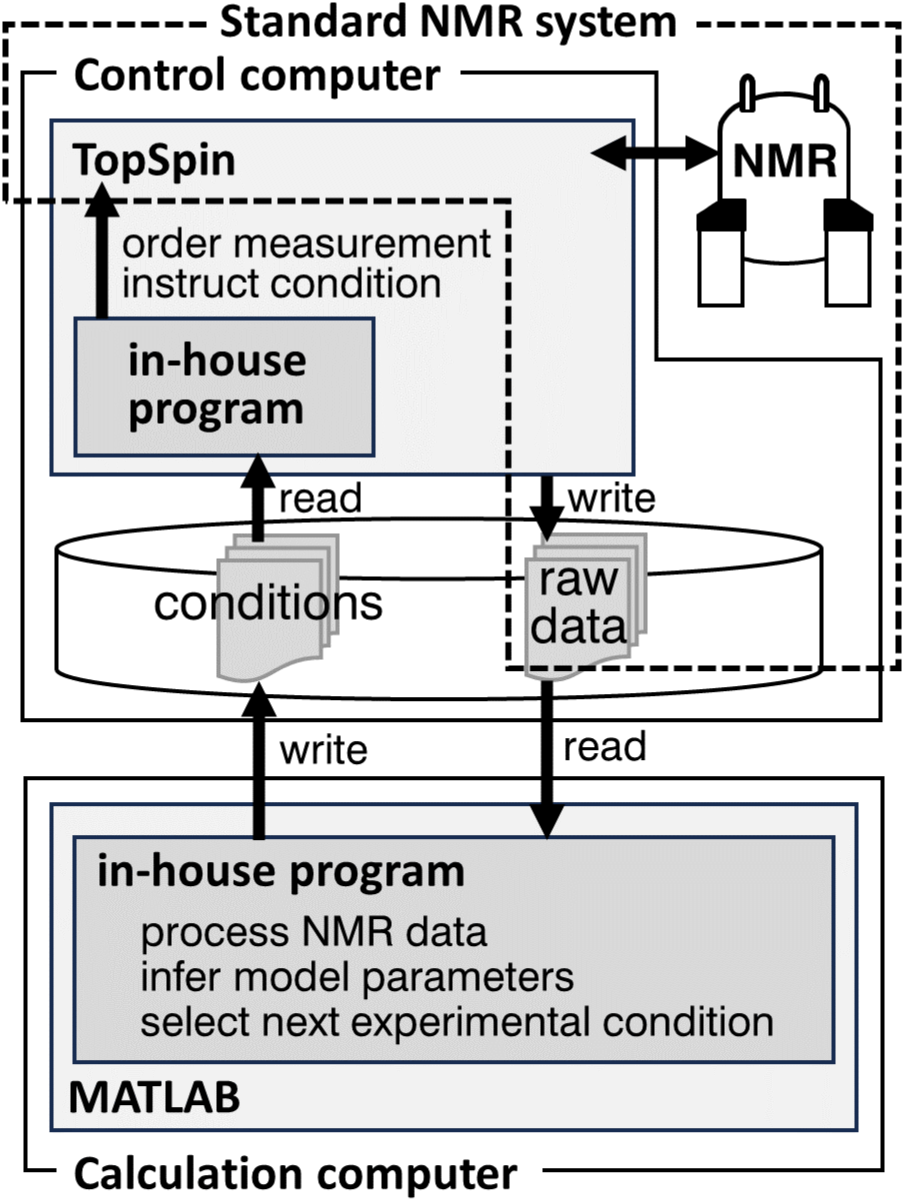
Schematic representation of the experimental configuration for adaptive CEST.

Fig 5b illustrates the model parameter estimates after 192 iterations of 2D measurements. The accuracy was evaluated with respect to the values of *p*_B_, *k*_ex_, and *ω*_B_ reported in the literature, separate inversion recovery or CPMG experiments for *R*_1_, *R*_2A_, and conventional CEST with 4-fold more scans for all parameters (Fig 5b). Adaptive CEST estimates agreed with these values. It outperformed, or at least performed comparable to, conventional CEST in terms of precision, except in the case of G39, which exhibited an outlying *ω*_B_ value.

## 3. General Discussion

In the preceding discussion, we assumed that the forward model and the noise were known. However, the proposed Bayesian experimental design method can also be implemented in general NMR experiments or even other applications by simply substituting the forward and noise models, while retaining calculation of mutual information using MCMC. As described in the Introduction, experimental designs pursuing the best inference of model parameters have been proposed in various research fields [9–16]. The method proposed in this paper serves as another example of a Bayesian optimal sequential experimental design, in which the computation algorithm does not depend on the type of measurement. Its applicability to NMR CEST measurements was validated by introducing an approximation for the forward model compatible with relatively heavy MCMC computations. One possible drawback of this method is its dependence on forward models. The proposed system considers a two-state forward model; thus, the system does not attempt to search for a third state after finding the second one. This is problematic in certain ^15^N-CEST applications involving models with more than two states; e.g., the observation of intermediate states in fold-unfold equilibria [37,38]. It should be noted that, in addition to general human-instructed research, it is possible to switch the model used for analysis after data acquisition. In this case, following adaptive measurement, all data are analyzed using a three-state model, expecting some information on the third state, despite the autonomous system not actively gathering such information. If the SNR is low, as assumed in this study, it is practical to prioritize identifying only one minor state over identifying all.

As discussed in the Theory section, it is usual in CEST to first analyze data on a per-residue basis to extract residue-specific *p*_B_ and *k*_ex_, and then switch to global fitting analysis assuming those values to be constant for all residues; the latter analysis is also applicable to adaptively acquired CEST data. However, the application of the proposed simple MCMC analysis to the global model is difficult because of the large dimensionality (5𝐾 + 2) of the parameter space.

To acquire responses corresponding to various frequency offsets efficiently, multifrequency or cosine-modulated saturation pulses have been used in some studies, instead of conventional single-frequency pulses [25–27]. The proposed method is capable of integrating these conventional and advanced pulses by considering them as different experimental condition candidates, provided that the forward model computation for advanced pulses is compatible with the MCMC calculations. The performance of the integrated method is expected to be better than or at least comparable to that of adaptive CEST with a conventional pulse.

In this study, the 2D measurement time varied with 𝑇_EX_. As we did not consider this variation over a single iteration, the objective of the experimental design was the amount of information gathered per iteration, not per unit time. If the measurement time is considered, the mutual information per unit measurement time may be a more suitable utility function. In this study, the NMR spectrometer was idle between measurements. It is worth considering to acquire data during this period as well, e.g., using 1D ^15^N-HSQC to monitor the state of the sample.

As discussed previously, in some situations, extending the iterations with different utility functions is useful. If, upon completion, an experiment yields insufficient general precision for model parameter estimates, more iterations can be added after the adaptive experiment using the same utility function. In this study, we performed the experiments with a fixed number of iterations; in this context, autonomous halting or extension of the experiment based on the required precision is an interesting topic.

Furthermore, we considered adaptive CEST with four different *ω*_1_ values and two different 𝑇_EX_ values. We did not attempt to wield more precise control over these parameters because the computation time for mutual information in the current algorithm was proportional to the number of discrete experimental conditions. In future works, a suitable experimental configuration should be identified for adaptive CEST.

## 4. Conclusions

The primary advantage of the adaptive optimization methods proposed in this paper is that it can select effective experimental conditions for model parameter estimation without human intervention, provided that a reliable forward model is available. In CEST, one of the model parameters (*ω*_B_) is almost unknown before starting the experiment, even though the design of the other parameters depends on it. In such cases, autonomous experimental design is helpful because of the independence of external instruction that it provides while switching the target model parameters. Moreover, for complicated nonlinear model functions, manual estimation of the current knowledge based on in-hand data, accurate prediction of the behavior of the observation with respect to varying model parameters, and appropriate experimental design to maximize information gathered about model parameters are difficult.

## 5. Materials and Methods

### 5.1 Sample preparation

The gene encoding the FF domain of HYPA/FBP11 containing the A39G mutation was synthesized by Eurofins Genomics K.K. (Tokyo, Japan). A template DNA encoding FF A39G with a histidine affinity tag and a Tobacco Etch Virus (TEV) protease cleavage site at its N-terminus was constructed using a polymerase chain reaction (PCR)-based method, as described previously [39]. U-^13^C/^15^N-labeled FF A39G was produced using the dialysis mode of a cell-free protein synthesis system, with a 9-mL inner reaction solution [40–43]. Affinity purification was performed by loading the reaction solution diluted with buffer A (20 mM Tris-Cl pH 8.0, 500 mM NaCl, and 20 mM imidazole) onto a HisTrap 1-mL affinity column (Global Life Sciences Solutions, Marlborough, MA, USA) and eluting with buffer B (identical to A, except containing 500 mM imidazole). The eluate was exchanged with buffer A via ultrafiltration with Amicon Ultra 15 MWCO = 3K (Merck, Darmstadt, Germany) and subsequently cleaved with 100 ug of TEV for 16 hours at 25 °C. The cleaved product was further purified using a HiTrap SP 1-mL cation exchange column (Global Life Sciences Solutions, Marlborough, MA, USA) with a gradient of C (50 mM 2-(N-morpholino)ethanesulfonic acid (MES)-sodium, pH 6.0) and D (same as C, except containing 1 M sodium chloride) buffers. Fractions containing FF A39G were concentrated and exchanged in the NMR buffer (100 mM sodium acetate, 200 mM sodium chloride, pH = 5.7) with Amicon Ultra 15 MWCO = 3 K (Merck, Darmstadt, Germany). The cleaved protein product was 71-residues long with no cloning artifacts at either end (GSQPAKKTYT WNTKEEAKQA FKELLKEKRV PSNASWEQGM KMIINDPRYS ALAKLSEKKQ AFNAYKVQTE K).

### 5.2 NMR measurements

All NMR measurements were performed at 274.2 K using an AVANCE III HD 700 MHz spectrometer equipped with a TCI CryoProbe (Bruker BioSpin, Rheinstetten, Germany). Prior to recording the NMR measurements, the sample temperature was calibrated with methanol following the manufacturer’s instructions. The B_1_ field strength was calibrated with a ^15^N-labeled Tryptophan and Glutamine solution using a pulse program “hsqc_cest_f3gpphtc_b1cal” provided by the manufacturer. For the CEST measurements, 500 uL of 0.1 mM U-^13^C/^15^N FF A39G in the NMR buffer was loaded into a 5 mm NMR sample tube. Main-chain amide signals were sequentially assigned to standard triple resonance experiments [44–46]. To verify the accuracy of CEST-derived relaxation rate constants, ^15^N R_1_ and R_2_ were separately measured with the same sample using pulse programs “hsqct1etf3gpsi3d” and “hsqct2etf3gpsi,” respectively. The spectra were processed and analyzed using MATLAB 2020a (MathWorks, MA, USA).

### 5.3 Adaptive CEST experiments

All CEST measurements were performed using a modified pulse program based on “hsqc_cest_etf3gpsitc3d”, which was originally designed for pseudo-3D measurement for ^1^H, ^15^N, and the irradiation-condition dimensions. We modified it to acquire a ^1^H-^15^N 2D plane under single irradiation conditions. Each 2D plane was acquired with 64 real points, 25-Hz spectral width, and 118.5-ppm carrier frequency in the ^15^N dimension, with eight cumulative transients corresponding to each time point. The adaptive NMR measurements were performed with the cooperation of two computers (Fig 6)—a control computer of the NMR system and a calculation computer (HP Z840 Workstation (HP Japan Inc., Tokyo, Japan) with dual Intel Xeon E5-2690v4 CPUs and 128 GiB main memory). On the control computer, an in-house C program running on TopSpin 3.5 software (Bruker BioSpin, Rheinstetten, Germany) was used to read the offset frequency, duration, and strength of each irradiation pulse from a local text file, start a measurement, and store raw time-domain data in local storage. On the calculation computer, another in-house program running on MATLAB 2020a (MathWorks, MA, USA) was used to write the irradiation conditions to the text file on the control computer, download the time-domain raw data, process them into spectra, analyze their peak intensities, perform MCMC for Bayesian inference of model parameters, and finally select each subsequent experimental condition based on mutual information. The in-house program utilized JSch 0.1.55 (JCraft, Sendai, Japan) for the SSH connection between the control and calculation computers, NMRglue 0.7 for NMR data processing [47], and mcmcstat for the MCMC analysis [48,49]. By the MCMC analysis, the following seven model parameters were inferred with either linear or logarithmic uniform prior distributions in the designated range: *p*_B_ ∈ [0, 0.1] (linear), *k*_ex_ ∈ [5, 1000] s^―1^ (logarithmic), *ω*_B_ ∈ [ ―1000, 1000] Hz (linear),*R*_1_ ∈ [0.1, 10] s^―1^ (logarithmic), *R*_2A_/*R*_1_ ∈ [1, 100] (logarithmic), **R**_2B_/*R*_1_ ∈ [1, 1000] (logarithmic), 𝐼_0_ ∈ [0.1, 10000] a.u. (logarithmic). The numbers of burn-in steps, sampling steps, and thinning intervals were 20,000, 30,000, and 50, respectively, in the mutual information calculation to reduce the computation time between NMR measurements. After all the iterations were completed, another iteration of MCMC with an increased number of steps (100,000 burn-in steps and 1,000,000 sampling steps) was performed for a more detailed analysis.

## Acknowledgements

The authors would like to express their earnest gratitude to the members of the Laboratory for Cellular Structural Biology for valuable discussions. We are grateful to S. Onami, principal investigator of the Research DX Foundation Team, for his understanding and support in publishing this work. We would also like to thank S. Yasuda, A. Yokooku, and A. Sekiguchi for their secretarial assistance.

## References

1. Box GEP, editor. Choice of response surface design and alphabetic Optimality1982.

2. Shannon CE. A mathematical theory of communication. Bell Syst Tech J. 1948;27: 379–423. doi: 10.1002/j.1538-7305.1948.tb01338.x.

3. Lindley DV. On a measure of the information provided by an experiment. Ann Math Statist. 1956;27: 986–1005. doi: 10.1214/aoms/1177728069.

4. Kullback S, Leibler RA. On information and sufficiency. Ann Math Statist. 1951;22: 79–86. doi: 10.1214/aoms/1177729694.

5. Cover TM, Thomas JA. Elements of information theory. Hoboken, New Jersey: John Wiley & Sons, Inc.; 1991.

6. Chaloner K, Verdinelli I. Bayesian experimental design: A review. Stat Sci. 1995;10: 273–304. doi: 10.1214/ss/1177009939.

7. Dror HA, Steinberg DM. Sequential experimental designs for generalized linear models. J Am Stat Assoc. 2008;103: 288–298. doi: 10.1198/016214507000001346.

8. Golovin D, Krause A. Adaptive submodularity: Theory and applications in active learning and stochastic optimization. J Artif Int Res. 2011;42: 427–486.

9. Cavagnaro DR, Myung JI, Pitt MA, Kujala JV. Adaptive design optimization: A mutual information-based approach to model discrimination in cognitive science. Neural Comput. 2010;22: 887–905. doi: 10.1162/neco.2009.02-09-959.

10. Vanlier J, Tiemann CA, Hilbers PAJ, van Riel NAW. A Bayesian approach to targeted experiment design. Bioinformatics. 2012;28: 1136–1142. doi: 10.1093/bioinformatics/bts092.

11. Drovandi CC, McGree JM, Pettitt AN. Sequential Monte Carlo for Bayesian sequentially designed experiments for discrete data. Comp Stat Data Anal. 2013;57: 320–335. doi: 10.1016/j.csda.2012.05.014.

12. Aggarwal R, Demkowicz MJ, Marzouk YM. Information-driven experimental design in materials science. In: Lookman T, Alexander FJ, Rajan K, editors. Information science for materials discovery and design. Cham: Springer International Publishing; 2016. pp. 13–44. doi: 10.1007/978-3-319-23871-5_2.

13. Ryan EG, Drovandi CC, McGree JM, Pettitt AN. A review of modern computational algorithms for Bayesian optimal design. Int Stat Rev. 2016;84: 128–154. doi: 10.1111/insr.12107.

14. Jiang H, Zhao Y. A review of Bayesian optimal experimental design on different models. In: Zhao Y, Chen D-G, editors. Modern statistical methods for Health Research. Cham: Springer International Publishing; 2021. pp. 205–220. doi: 10.1007/978-3-030-72437-5_10.

15. Kalinin SV, Ziatdinov M, Hinkle J, Jesse S, Ghosh A, Kelley KP, et al. Automated and autonomous experiments in electron and scanning probe microscopy. ACS Nano. 2021;15: 12604–12627. doi: 10.1021/acsnano.1c02104.

16. Lee SY. Bayesian nonlinear models for repeated measurement data: An overview, implementation, and applications. Mathematics. 2022;10: 898. doi: 10.3390/math10060898.

17. Eghbalnia HR, Bahrami A, Tonelli M, Hallenga K, Markley JL. High-resolution iterative frequency identification for NMR as a general strategy for multidimensional data collection. J Am Chem Soc. 2005;127: 12528–12536. doi: 10.1021/ja052120i.

18. Bahrami A, Tonelli M, Sahu SC, Singarapu KK, Eghbalnia HR, Markley JL. Robust, integrated computational control of NMR experiments to achieve optimal assignment by ADAPT-NMR. PLOS ONE. 2012;7: e33173. doi: 10.1371/journal.pone.0033173.

19. Song YQ, Tang Y, Hürlimann MD, Cory DG. Real-time optimization of nuclear magnetic resonance experiments. J Magn Reson. 2018;289: 72–78. doi: 10.1016/j.jmr.2018.02.009.

20. Waudby CA, Burridge C, Christodoulou J. Optimal design of adaptively sampled NMR experiments for measurement of methyl group dynamics with application to a ribosome-nascent chain complex. J Magn Reson. 2021;326: 106937. doi: 10.1016/j.jmr.2021.106937.

21. Vallurupalli P, Sekhar A, Yuwen T, Kay LE. Probing conformational dynamics in biomolecules via chemical exchange saturation transfer: A primer. J Biomol NMR. 2017;67: 243–271. doi: 10.1007/s10858-017-0099-4.

22. Vallurupalli P, Bouvignies G, Kay LE. Studying “invisible” excited protein states in slow exchange with a major state conformation. J Am Chem Soc. 2012;134: 8148–8161. doi: 10.1021/ja3001419.

23. Bolik-Coulon N, Hansen DF, Kay LE. Optimizing frequency sampling in CEST experiments. J Biomol NMR. 2022;76: 167–183. doi: 10.1007/s10858-022-00403-2.

24. Carneiro MG, Reddy JG, Griesinger C, Lee D. Speeding-up exchange-mediated saturation transfer experiments by Fourier transform. J Biomol NMR. 2015;63: 237–244. doi: 10.1007/s10858-015-9985-9.

25. Leninger M, Marsiglia WM, Jerschow A, Traaseth NJ. Multiple frequency saturation pulses reduce CEST acquisition time for quantifying conformational exchange in biomolecules. J Biomol NMR. 2018;71: 19–30. doi: 10.1007/s10858-018-0186-1.

26. Yuwen T, Bouvignies G, Kay LE. Exploring methods to expedite the recording of CEST datasets using selective pulse excitation. J Magn Reson. 2018;292: 1–7. doi: 10.1016/j.jmr.2018.04.013.

27. Yuwen T, Kay LE, Bouvignies G. Dramatic decrease in CEST measurement times using multi-site excitation. ChemPhysChem. 2018;19: 1707–1710. doi: 10.1002/cphc.201800249.

28. Ceccon A, Schmidt T, Tugarinov V, Kotler SA, Schwieters CD, Clore GM. Interaction of huntingtin Exon-1 peptides with lipid-based micellar nanoparticles probed by solution NMR and Q-band pulsed EPR. J Am Chem Soc. 2018;140: 6199–6202. doi: 10.1021/jacs.8b02619.

29. Kragelj J, Orand T, Delaforge E, Tengo L, Blackledge M, Palencia A, et al. Enthalpy– entropy compensation in the promiscuous interaction of an intrinsically disordered protein with homologous protein partners. Biomolecules. 2021;11: 1204. doi: 10.3390/biom11081204.

30. Maciejewski MW, Mobli M, Schuyler AD, Stern AS, Hoch JC. Data sampling in multidimensional NMR: Fundamentals and strategies. In: Billeter M, Orekhov V, editors. Novel sampling approaches in higher dimensional NMR. Berlin, Heidelberg: Springer Berlin Heidelberg; 2012. pp. 49–77. doi: 10.1007/128_2011_185.

31. Palmer AG. Chemical exchange in biomacromolecules: Past, present, and future. J Magn Reson. 2014;241: 3–17. doi: 10.1016/j.jmr.2014.01.008.

32. McConnell HM. Reaction rates by nuclear magnetic resonance. J Chem Phys. 1958;28: 430–431. doi: 10.1063/1.1744152.

33. Trott O, Palmer AG. R_1ρ_ relaxation outside of the fast-exchange limit. J Magn Reson. 2002;154: 157–160. doi: 10.1006/jmre.2001.2466.

34. Baldwin AJ, Kay LE. An R_1ρ_ expression for a spin in chemical exchange between two sites with unequal transverse relaxation rates. J Biomol NMR. 2013;55: 211–218. doi: 10.1007/s10858-012-9694-6.

35. Houlsby N, Huszár F, Ghahramani Z, Lengyel M. Bayesian active learning for classification and preference Learning2011; 2011:[arXiv:1112.5745 p.]. Available from: https://ui.adsabs.harvard.edu/abs/2011arXiv1112.5745H.

36. Nagashima K. B_1_ mapping of liquid NMR transversal RF coils: Analysis of heterogeneity and nonlinearity. Concepts Magn Reson B. 2012;41B: 1–12. doi: 10.1002/cmr.b.20211.

37. Lim J, Xiao T, Fan J, Yang D. An off-pathway folding intermediate of an acyl carrier protein domain coexists with the folded and unfolded states under native conditions. Angew Chem Int Ed Engl. 2014;53: 2358–2361. doi: 10.1002/anie.201308512.

38. Tiwari VP, Toyama Y, De D, Kay LE, Vallurupalli P. The A39G FF domain folds on a volcano-shaped free energy surface via separate pathways. Proc Natl Acad Sci U S A. 2021;118: e2115113118. doi: 10.1073/pnas.2115113118.

39. Yabuki T, Motoda Y, Hanada K, Nunokawa E, Saito M, Seki E, et al. A robust two-step PCR method of template DNA production for high-throughput cell-free protein synthesis. J Struct Funct Genomics. 2007;8: 173–191. doi: 10.1007/s10969-007-9038-z.

40. Kigawa T. Cell-free protein production system with the *E. coli* crude extract for determination of protein folds. In: Endo Y, Takai K, Ueda T, editors. Cell-free protein production. Methods and protocols. Totowa, New Jersey: Humana Press; 2010. pp. 101–111. doi: 10.1007/978-1-60327-331-2_10.

41. Kigawa T, Muto Y, Yokoyama S. Cell-free synthesis and amino acid-selective stable isotope labeling of proteins for NMR analysis. J Biomol NMR. 1995;6: 129–134. doi: 10.1007/BF00211776.

42. Kigawa T, Yabuki T, Matsuda N, Matsuda T, Nakajima R, Tanaka A, et al. Preparation of *Escherichia coli* cell extract for highly productive cell-free protein expression. J Struct Funct Genomics. 2004;5: 63–68. doi: 10.1023/B:JSFG.0000029204.57846.7d.

43. Matsuda T, Koshiba S, Tochio N, Seki E, Iwasaki N, Yabuki T, et al. Improving cell-free protein synthesis for stable-isotope labeling. J Biomol NMR. 2007;37: 225–229. doi: 10.1007/s10858-006-9127-5.

44. Bax A, Grzesiek S. Methodological advances in protein NMR. Acc Chem Res. 1993;26: 131–138. doi: 10.1021/ar00028a001.

45. Clubb RT, Thanabal V, Wagner G. A constant-time three-dimensional triple-resonance pulse scheme to correlate intraresidue ^1^H^N^, ^15^N, and ^13^C’ chemical shifts in ^15^N·^13^C-labelled proteins. J Magn Reson (1969). 1992;97: 213–217. doi: 10.1016/0022-2364(92)90252-3.

46. Ikura M, Kay LE, Bax A. A novel approach for sequential assignment of ^1^H, ^13^C, and ^15^N spectra of larger proteins: Heteronuclear triple-resonance three-dimensional NMR spectroscopy. Application to calmodulin. Biochemistry. 1990;29: 4659–4667. doi: 10.1021/bi00471a022.

47. Helmus JJ, Jaroniec CP. Nmrglue: An open source Python package for the analysis of multidimensional NMR data. J Biomol NMR. 2013;55: 355–367. doi: 10.1007/s10858-013-9718-x.

48. Haario H, Laine M, Mira A, Saksman E. DRAM: Efficient adaptive MCMC. Stat Comput. 2006;16: 339–354. doi: 10.1007/s11222-006-9438-0.

49. Haario H, Saksman E, Tamminen J. An adaptive metropolis algorithm. Bernoulli. 2001;7: 223–242. doi: 10.2307/3318737.

